# Anionic polymers stabilize Cas9 ribonucleoprotein nanoparticles to improve direct protein delivery and genome editing efficiencies in plant protoplasts

**DOI:** 10.1101/2025.02.02.635105

**Authors:** Joshua D. Hubbard, Sophia Tomatz, Nicole Carll, Juliana Matos, Danya Hassan, Tong Ly, Markita P. Landry

## Abstract

The application of CRISPR-based genome editing tools in plants is often challenged by low editing efficiencies, requiring most plant editing workflows to proceed through the delivery of pre-formed ribonucleoprotein (RNP) complexes to protoplasts. Here, we report increases in protoplast-based RNP delivery and genome editing efficiencies through the addition of anionic polymers to standard protoplast transfection protocols. We test addition of various polymers and peptides for their ability to increase genome editing efficiencies in both *Nicotiana benthamiana* and *Arabidopsis thaliana* protoplasts, by adding these components to standard PEG-based protoplast transfection protocols: *i)* non-covalent addition of charged polymers, *ii)* non-covalent addition of amphiphilic peptide A5K, and *iii)* tyrosinase-mediated covalent conjugation of various relevant peptide motifs directly to the RNP. Incorporation of the amphiphilic peptide A5K or covalent attachment of peptides to the RNP had no positive effect on editing efficiencies. However, we found that addition of anionic polymer polyglutamic acid to standard PEG transfection protocols significantly improved editing efficiencies in both *Nicotiana benthamiana* and *Arabidopsis thaliana* protoplasts relative to RNPs alone without negatively impacting protoplast viability. Our results suggest anionic polymers stabilize the RNP and increase the colloidal stability of the protoplast transfection workflow. This simple and straightforward method of stabilizing Cas9 RNPs can be easily adopted by others working on direct protein delivery to plant protoplasts to increase genome editing efficiencies.

**Key message:** The addition of anionic polymer polyglutamic acid to standard protoplast PEG transfection workflows enhance CRISPR-Cas9-mediated gene editing in plant protoplasts. Our results suggest the mechanism of increased transfection efficiency is due to colloidal stabilization of RNPs.

**Graphical Abstract:** 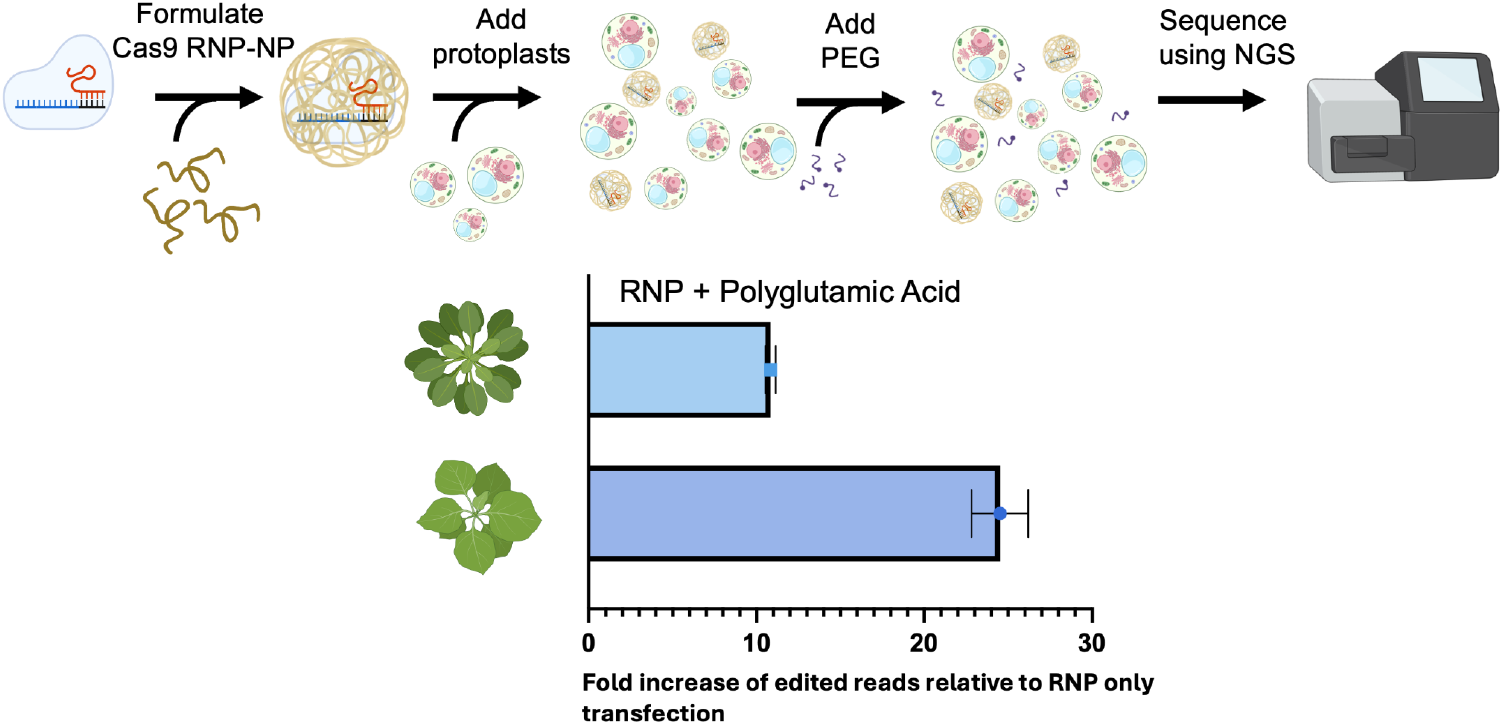

## Introduction

The emergence of CRISPR/Cas-based biotechnologies over a decade ago precipitated an expansive and persistent revolution in the field of plant genetic engineering. However, there remains a need to improve on methods to efficiently deliver CRISPR reagents, particularly in plant systems where genome editing efficiencies remain low relative to non-plant counterparts. For genome editing applications in agriculture, direct delivery of the guide RNA and Cas protein preassembled into a ribonucleoprotein (RNP) is of particular interest, as editing does not require transgene insertion, thus circumventing the need to cross out transgenes and assuaging the regulatory process of adopting edited crops into the marketplace (Landry and Mitter 2019). Furthermore, RNPs are only transiently present before being degraded by endogenous enzymes, compared to constitutive Cas9/gRNA expression, resulting in fewer off-target effects and reduced overall cell toxicity than with nucleic acid-based transfection (Zhang et al. 2021). RNP delivery in mammalian systems has been well-established in a range of cell types using a variety of delivery approaches, including successful demonstrations of advanced base-editors and guide multiplexing (Anzalone et al. 2020). In plants, RNP delivery of three of the basic CRISPR systems–Cas9, Cas12a, Cas12b–have been the focus of most studies, while only recently have other variants (*e*.*g*. Casϕ) and advanced base- and prime-editors begun to be explored (Zhang et al. 2021; Wang et al. 2021; Li et al. 2023). The primary distinguishing feature of plant cells in the context of direct delivery of proteins is the cell wall, which functions to exclude exogenous materials. There is a hypothesized size exclusion limit of 5-10 nm, which corresponds to a protein size exclusion limit of 50-100 kDa (Squire et al. 2023). To circumvent the plant cell wall obstacle, most protein delivery in plants is accomplished in protoplasts–plant cells that have had the cell wall enzymatically degraded (Jiang et al. 2021). However, even in the absence of the cell wall, protoplast genome editing efficiencies overall are highly variable across and within species, with many studies reporting editing efficiencies as low as 0.1%, and with editing efficiencies of over 25% only reported in several species (Zhang et al. 2021; Wang et al. 2021). Delivery of RNPs to mammalian cell systems is routinely much higher (>50%) than in plants (Liang et al. 2015), suggesting the presence of additional, unexplained barriers to CRISPR-mediated gene editing in plants.

It has previously been reported that the inclusion of non-homologous oligonucleotides or additional guide RNA can increase editing efficiencies of RNP electroporation in mammalian cells (Richardson et al. 2016; Roth et al. 2018). These findings motivated our investigation of non-oligonucleotide polymers and peptides that may harness this same unknown mechanism for improved transfection outcomes. Notably, prior work has found that incorporation of commercially synthesized anionic polymers into the RNP complex showed success in both improving gene editing efficiencies and subsequent survival rates after electroporation-mediated Cas9 transfection of mammalian cells, while cationic polymers had a detrimental effect (Nguyen et al. 2020). In contrast to the hypothesized mechanism of action associated with non-homologous oligonucleotide-based methods–which suggest these oligos elicit a cellular response that impacts double-stranded break (DSB) repair–anionic polymers increase editing due to improved colloidal stability of the RNP complex provided by anionic polymers prior to, and during, delivery. A proposed shared mechanism of action for the stabilizing effects of anionic polymers and nucleotides is the shielding of positively charged residues of the Cas9 protein from exposed negatively charged portions of Cas9-bound gRNA, thus improving colloidal stability. This stability is hypothesized to be achieved through the electrostatic association of the anionic polymer or oligonucleotide with positively charged residues exposed on the surface of the RNP, preventing aggregation (Nguyen et al. 2020). Given this proposed mechanism can be reasonably assumed to be cell-type independent, we reasoned that the inclusion of anionic polymers may have a similar effect in protoplast transfection.

In plants, protein delivery and, more specifically, genome editing nuclease delivery, is predominantly accomplished through PEG-mediated transfection of protoplasts or biolistic delivery to plant tissues (Jiang et al. 2021; Zhang et al. 2021; Wang et al. 2021). Both PEG-mediated and biolistic delivery of genome editing nucleases suffer from low editing efficiencies that challenge the use of RNPs in plant biotechnology. Recently-developed methods for quantitative measurement of protein delivery efficiency have been used to demonstrate that covalent attachment of cell penetrating peptides to small protein cargos can enable uptake into intact plant tissues (Wang et al. 2023). However, generating chimeric peptide-protein constructs can often be time-, labor-, and resource-intensive, and genome editing protein cargoes may be too large to benefit from cell penetrating peptide-based delivery. In addition, chimeric products are generally less stable, and, due to their cationic nature, more toxic to cells, limiting their scope of application (Wang et al. 2021). Certain classes of peptides, such as those associated with endosome disruption, are incompatible with traditional covalent synthesis methods due to their insolubility (Reissmann 2014). With this context, it is clear that straightforward methods to improve genome editing efficiencies in plants remain sparse.

In this work, we investigate the ability of a library of polymers and peptides to stabilize RNPs to increase Cas9 gene editing efficiencies in plant protoplasts. To confirm the colloidal stability of RNP complexes formed with these various peptides and polymers, we first screen their ability to generate DNA cutting *in vitro*. Next, we test the effect of the polymers on *in vivo* genome editing efficiencies in two different protoplast species. We find that the inclusion of polyglutamic acid (PGA) polymers during sgRNA-Cas9 complex formation improves gene editing rates in protoplasts derived from both *Nicotiana benthamiana* and *Arabidopsis thaliana* plants without affecting protoplast viability post-transfection. We further show that PGA is unique in its ability to increase protoplast genome editing efficiencies, with no increases in genome editing efficiencies observed for other polymers and peptides tested. We propose that PGA might be more broadly used for protoplast transfection experiments, particularly for those seeking to deliver genome editing nucleases and enhance genome editing efficiencies.

## Results

### Polymer-stabilized Cas9 RNPs exhibit improved gene editing in vitro

We generated a small library of 10 polymers and peptides to screen for their ability to improve editing efficiencies following their addition to standard PEG-mediated protoplast transfection (Fig. 1a). This library included five cationic and anionic polymers based on their prior use for Cas9-based delivery and transfection in mammalian systems: poly-L-glutamic acid (PGA), poly-L-aspartic acid (PLD), polyacrylic acid (PAA), poly-L-arginine (PLR), and protamine sulfate (PS) (Nguyen et al. 2020). An additional five peptides previously identified as potentially exhibiting cell-penetrating behavior in mammalian and plant systems were also tested: AK5, FLG22, SV40, WUSCHEL (WUS), and Nona-L-arginine (R9) (Fig. 1a) (Robatzek et al. 2006; Yadav et al. 2011; Lobba et al. 2020; Foss et al. 2023; Wang et al. 2023). As the mechanism of RNP uptake to protoplasts remains unclear, we screened peptides that have cell penetrating activity, NLS motifs, or unique functions in plants. Our polymer library included cationic and anionic polymers chosen from a prior screen in mammalian cell systems, with specific anionic polymers implicated in increasing RNP stability, thus generating enhanced editing outcomes (Nguyen et al. 2020). We also tested amphiphilic peptide A5K, due to its prior use in peptide-enabled RNP delivery for CRISPR engineering (PERC method). A5K contains an endosomolytic domain and a cationic cell penetrating domain, which, when incorporated into the RNP complex, improves editing outcomes (Foss et al. 2023). The cationic domain is hypothesized to interact electrostatically with the negatively-charged RNP surface while displaying the endosomolytic domain.

**Fig. 1.**
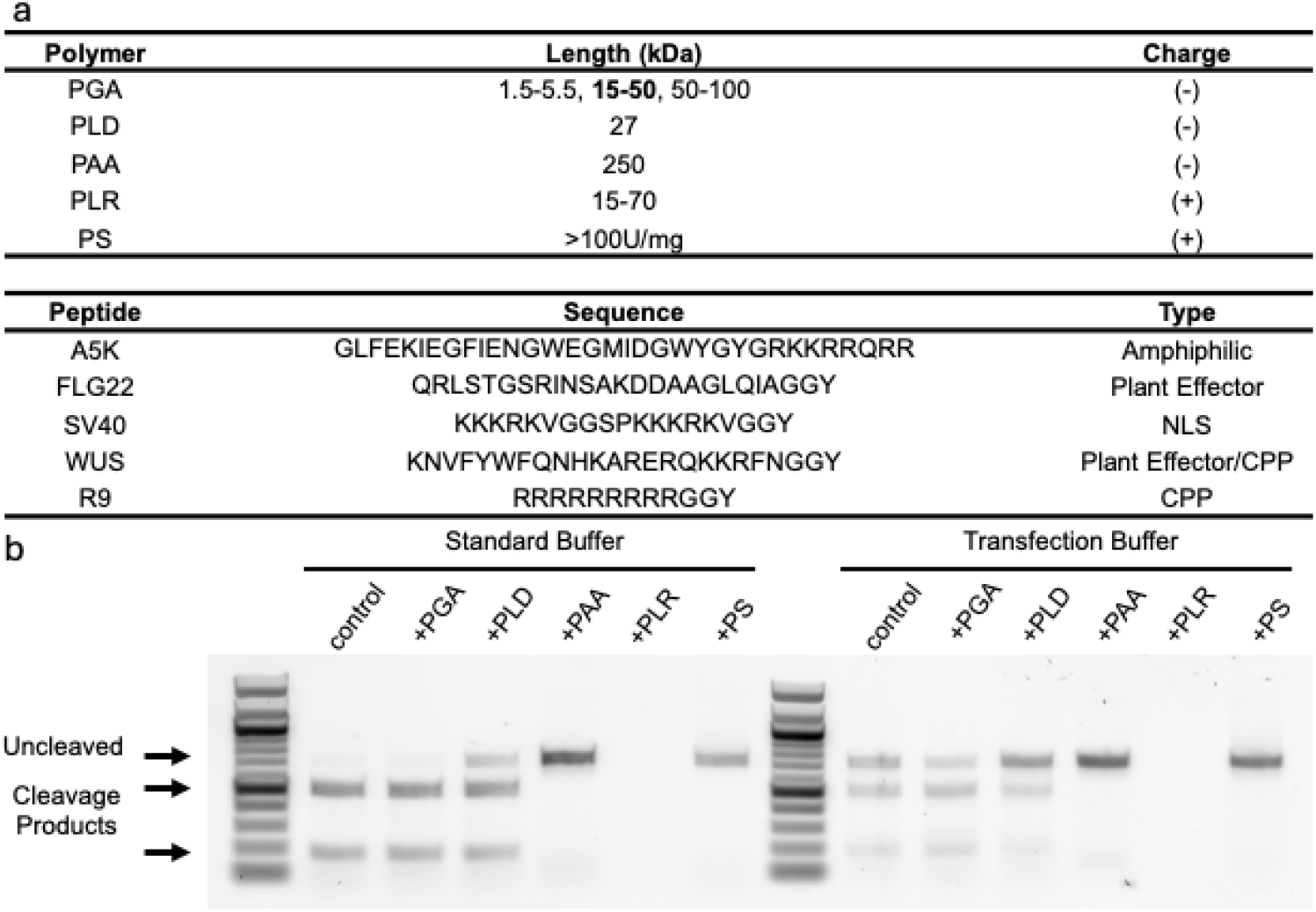
**(a**) (top) Anionic and cationic polymers of various lengths employed in this study. (bottom) Functional peptide motifs screened in this study were identified from previously published literature. **(b)** *In vitro* cleavage assay is used to screen for enzymatic activity of Polymer-RNP NPs. The anionic polymers PGA and PLD were the only polymers that exhibited cleavage activity. PAA and PS show uncleaved bands, while PLR inhibited the DNA from running on the gel.

Finally, we explored whether covalent conjugation of chemical aides to the Cas9 RNP could be used to improve RNP-mediated editing. In mammalian systems, the direct linkage of functional peptides to Cas9 proteins to improve RNP delivery and gene editing has been explored through genetic fusions and chemical conjugation methods (Staahl et al. 2017; Lobba et al. 2020). For these experiments, we used a small library of peptides previously tested in mammalian systems as well as novel, plant derived peptides. Peptides tested in mammalian systems included SV40 and R9, which has also been tested in plant systems (Kosuge et al. 2008; Staahl et al. 2017; Numata et al. 2018; Wang et al. 2023). We also incorporated plant and plant-pathogen derived peptides implicated in endogenous cell trafficking and uptake pathways into our screen. WUSCHEL (WUS), a homeodomain transcription factor expressed in shoot meristematic cells, is known to migrate from its site of synthesis in the organizing center (OC) to the central zone (CZ) (Yadav et al. 2011). The plant pathogen-derived conserved peptide FLG22 was chosen due to its interaction with the plant receptor FLS2, which is known to induce receptor mediated endocytosis into plant cells (Robatzek et al. 2006). The latter four peptides (SV40, R9, WUS, FLG22) all have a GGY motif included at the C-terminus to allow for tyrosinase-mediated conjugation to the surface exposed cysteines on Cas9.

Having finalized our polymer library as justified above, we next performed *in vitro* DNA cleavage assays to confirm the ability of RNPs to cut their DNA target sites in the presence of each polymer. We found that the cationic polymers poly(L-arginine) (PLR) and protamine sulfate (PS) abolished enzymatic activity (Fig. 1b). RNPs formed in the presence of anionic polymers polyglutamic acid (PGA) and poly-L-arginine (PLD) both exhibited similar cleavage activity relative to the polymer-free control, with PGA generating the most complete cleavage. Notably, formulations including polyacrylic acid (PAA) did not exhibit any *in vitro* cleavage activity, despite polyacrylic acid being an anionic polymer (Fig. 1b). PGA demonstrated successful *in vitro* cleavage at previously reported concentrations and polymer lengths (SI Fig. 1) (Nguyen et al. 2020). These results, along with results from mammalian systems indicating improved editing with PGA, led us to further explore the *in vivo* activity of PGA-RNPs in plant protoplasts.

### Polymer-stabilized Cas9 RNPs exhibit improved gene editing in protoplasts

We next tested whether the inclusion of anionic polymers increased gene editing efficiency of PEG-mediated protoplast transfection with RNPs. Anionic polymers, sgRNA, and apo-spCas9 and were combined to a final concentration of 5mg/mL, 6μM, and 4μM respectively. We used chemically synthesized sgRNAs designed based on protospacer sequences that had previously been identified as successfully targeting the *AtPDS3* gene in *A. thaliana* (Li et al. 2013) and the *NbPDS* gene in *N. benthamiana* (Nekrasov et al. 2013). These PDS target genes encode for phytoene desaturase enzymes that, when disrupted, results in albino phenotypes and dwarfed seedlings due to their role in carotenoid biosynthesis (Qin et al. 2007). Deep amplicon sequencing was used to identify insertions and deletions in the targeted regions of the genome. In *Nicotiana benthamiana* protoplasts, the addition of PGA to the RNP complex increased gene editing efficiency from 0.44±0.15 % in a PEG-only RNP delivery condition, to 10.8±1.7 %, representing a roughly 25-fold increase due to the addition of PGA (Fig. 2a). Similarly, in *Arabidopsis thaliana*, the addition of PGA to the RNP complex increased gene editing efficiency from 0.19±0.06 % to 2.1±0.5 which is a 10.8±0.30-fold increase relative to PEG-only RNP delivery (Fig. 2b). The viability of *A. thaliana* protoplasts 24 hr post-transfection was not negatively impacted by the inclusion of PGA, although no protective effect was observed either (Fig. 2b). It is worth noting the PEG transfection itself does reduce cell viability, which might limit overall editing efficiencies, but PEG-related viability losses are consistent across RNP treatments. Additional experiments in *N. benthamiana* revealed that increasing the concentration of PGA 2- and 5-fold relative to the standard concentration of PGA reduced editing efficiency, and that editing efficiencies were higher for protoplasts incubated at 26°C vs. room temperature after transfection (SI Fig. 2). Protoplast experiments at temperatures above 26°C lead to evident decreases cell viability, leading to low or abolished editing. Other polymers tested at previously reported concentrations–poly(L-aspartic acid) (PLD) at 0.665 mg/mL, poly(acrylic acid) (PAA) at 3.6 mg/mL, poly(L-arginine) (PLR) at 5 mg/mL, and protamine sulfate (PS) at 0.5 mg/mL–resulted in a decrease or lack of in genome editing events relative to standard PEG transfection protocol, consistent with *in vitro* cleavage results (data not shown).

**Fig. 2.**
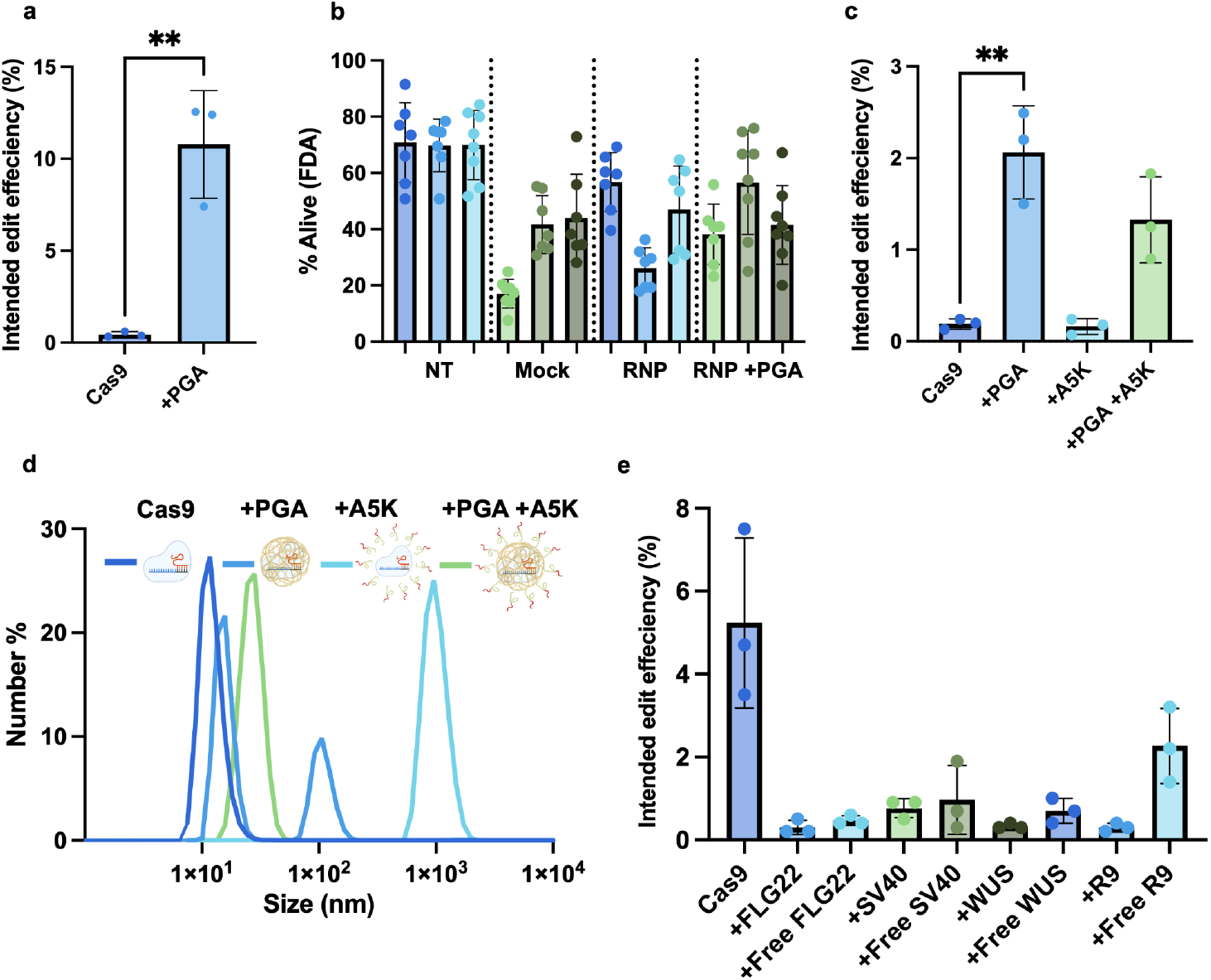
Gene editing efficiencies of PEG-transfected Cas9 RNPs to protoplasts that were **(a)** formulated with or without the anionic polymer PGA to *N. benthamiana* protoplasts. For all editing results, editing efficiencies were quantified using CRISPResso2 software to analyze NGS results. Mean and SD of three technical replicates are shown. Unpaired *t* test was used to calculate *P* value of indicated comparisons. \*\**P* < 0.01. **(b)** Viability of *A. thaliana* protoplasts pre- and post-transfection was evaluated using FDA staining. Data is shown for three individual wells for each condition, representing biological replicates of each treatment. The mean and SD of FDA positive cells are plotted for each well, with each data point representing the percentage of FDA positive cells in a single field of view in each well. Representative images are shown in SI Fig. 3. **(c)** Gene editing efficiencies of PEG-transfected Cas9 RNPs that were formulated with anionic polymer PGA alone, A5K alone, or both PGA and A5K to *A. thaliana* protoplasts. **(d)** The hydrodynamic radius of RNP-nanoparticles in NEBuffer3 was evaluated by DLS. **(e)** Cas9 RNP editing efficiencies in *N. benthamiana* protoplasts with peptides conjugated or unconjugated (free). Mean and SD of three technical replicates are shown.

### Non-covalently attached AK5 peptide (PERC method) does not improve editing efficiency in protoplasts

Different functional peptides have been employed to improve uptake of RNPs in recalcitrant mammalian cell types, with some notable success (Lobba et al. 2020; Foss et al. 2023). We first tested whether addition of the AK5 peptide would improve genome editing efficiency in *Arabidopsis* protoplasts when added to colloidal suspensions of RNPs prior to PEG transfection, referred to as free peptide addition. For A5K, the peptide was combined at a molar ratio of 10:1 peptide:RNP, and compared to editing achieved using plain RNPs and PGA-RNPs. We adapted the PERC method of preparing RNPs for PEG-mediated protoplast transfection, incubating 40 μM of the A5K peptide with the RNP complex for five minutes before addition of *Arabidopsis* protoplasts followed by PEG (Fig. 1a). Sequencing revealed no editing advantage of A5K relative to a PEG-only transfection protocol in protoplasts (Fig. 2c). Next, we sought to test if we could recover polymer-driven increases in genome editing efficiency by testing the synergistic effects of both A5K and PGA polymers together. We prepared RNPs for PEG-mediated protoplast transfection including 40 μM A5K and 5mg/mL PGA for a 5-min incubation prior to protoplast addition. We find that both A5K and PGA together yield a modest 6.95±0.30-fold increase in editing efficiency relative to the PEG-only control and together confer no advantage in genome editing efficiency relative to the addition of PGA alone (Fig. 2c).

Given the ability of PGA to improve editing and recover some of the cleavage activity lost due to A5K addition, we sought to characterize the physicochemical properties of RNPs in the presence of A5K and PGA polymers. Dynamic light scattering was employed to evaluate the stability of polymer- and peptide-based RNP-NPs. Incorporation of PGA with the RNP resulted in an increase in the measured hydrodynamic size of the protein, which is consistent with the hypothesis that the polymer is coating or otherwise complexing with the RNP (Fig. 2d). When A5K was introduced to RNPs in the absence of PGA, a large DLS peak emerged around 1×10^3^ nm, indicating the formation of an unstable precipitate which could be observed as a visibly cloudy solution. Alternatively, when A5K was introduced to Cas9 RNPs stabilized by PGA, the solution remained stable with a single peak around thirty nanometers (Fig 2c). These results suggest that PGA is able to re-stabilize a colloidally unstable RNP solution, further supporting the hypothesis that the gene editing increase observed with PGA polymer addition results from a mechanism involving stabilization of the RNP colloidal suspension. Considered in conjunction with gene editing results, these findings suggest that amphiphilic peptide-based transfection methods may be more consistently stable and successful at facilitating delivery of RNPs when employed in combination with anionic polymer stabilization, specifically through addition of PGA.

### Covalent attachment of peptides yields Cas9 RNPs that are not active in planta

Next, we sought to explore whether covalent peptide attachment could improve editing using peptides with exposed cysteine residues previously tested in mammalian systems and derived from plants and their pathogens. The inclusion of multiple nuclear localization signals (NLS) through genetic fusions to the N- and C-termini of Cas9 improved editing efficiencies and has become the standard in the field for most applications where transport across the nuclear envelope is necessitated (Staahl et al. 2017). Because genetic fusions as an approach are limited in their application, since this method only allows for peptides to be incorporated at surface exposed termini, researchers recently employed tyrosine-mediated coupling to facilitate post-translational and site-selective linkage of peptides to the two surface-exposed native cysteine residues at positions 80 and 574 of Cas9 (Lobba et al. 2020). This simple and straightforward approach allowed for the rapid screening of Cas9-peptide conjugates for enhanced gene editing, revealing that the incorporation of additional SV40 NLS tags enhanced gene editing in human neural progenitor cells (Lobba et al. 2020). In plants, only recently have genetic fusions of small proteins–but not yet genome editing nucleases–been shown to enable direct delivery of small proteins into plant cells (Wang et al. 2023). Specifically, peptide conjugation work in the plant space has focused on quantifying uptake of biomolecules across the plant cell wall. In particular, this work demonstrated that covalent conjugation of cell-penetrating peptide domains–including R9–was essential for uptake of small proteins and peptides to mature plant tissues (Wang et al. 2023). Finally, we introduced two novel peptides implicated in cell trafficking, WUS and FLG22.

Given this context, we were interested in whether the *covalent–*relative to the above-discussed colloidal–incorporation of cell penetrating peptide tags would enhance RNP delivery to *N. benthamiana* plant protoplasts, either past the cell membrane or the endosomal membrane after PEG-mediated cell uptake. We generated four Cas9-peptide conjugates to screen in protoplasts (Fig. 1a). Successful covalent attachment at a 2:1 molar ratio of peptide:protein was confirmed using ESI-TOF MS for each of the conjugates generated (SI Fig. 4). Each peptide was then tested for its ability to increase genome editing efficiency in PEG-mediated transfection of *N. benthamiana* protoplasts with the chimeric Cas9 protein, relative to unmodified Cas9, and unconjugated colloidal addition of the peptide. The RNP or RNP-peptide conjugates were at a concentration of 4 μM and, when incorporated, the peptides were added at 2:1 molar ratio of peptide:unmodified RNP. We used a different sgRNA guide to target the *NbPDS* gene of *N. benthamiana* taken from previously published literature (Uranga et al. 2023). Unlike their ability to improve genome editing efficiencies in mammalian transfection studies (Lobba et al. 2020) we found that none of the Cas9 chimeric proteins exhibited any editing events, relative to 5.2±2.1 % genome editing efficiencies measured with unmodified Cas9. Furthermore, we tested whether covalent versus noncovalent addition of the peptide resulted in better genome editing outcomes in a PEG transfection protocol. Interestingly, in most cases, the covalently attached Cas9-peptide conjugated underperformed relative to Cas9 mixed with unconjugated peptide at an equivalent concentration (Fig. 2e). These results suggest that covalent attachment of peptides may not be effective in the context of protoplast membrane translocation compared to uptake in mammalian systems (Lobba et al. 2020).

In the course of sequencing experiments for the above RNP-based methods, mock transfections were also conducted across different experiments wherein PEG free transfection buffer was used in place of PEG-containing buffer. However, PEG-free mock transfection conditions did not produce editing (data not shown), indicating the role of PEG in driving RNP uptake, even when RNPs are modified with cell-penetrating elements. However, further optimization of PEG-free methods should be done to evaluate the efficacy of these RNP formulations for cell transfection independent of PEG.

## Discussion

Our study of various peptides and polymer formulations adapted from mammalian systems revealed that colloidal stability is a significant factor impacting gene editing efficiencies of Cas9 RNP delivery to plant protoplasts. The inclusion of the commercially available anionic polymer PGA improved PEG-mediated protoplast gene editing efficiencies by 12.4±1.7-fold and 10.8±0.30-fold for *N. benthamiana* and *A. thaliana* respectively, relative to standard PEG transfection protocols. This finding is consistent with previously published results in mammalian systems, supporting the hypothesis that anionic polymer addition stabilizes the Cas9 RNPs across two different plant species tested. Cleavage results reinforce that a negative charge of the polymer is critical to retain enzymatic activity. Furthermore, DLS data showing an increase in the radius of gyration of the RNP-NP relative to the plain RNP supports the hypothesis that stabilization is achieved through a surface interaction between the RNP and the PGA polymer. While our results show promise for translating certain transfection aides used in mammalian cells to plant systems, this work showed that not all methods can be directly applied with similar success. Screening of peptides using both covalent and non-covalent approaches, both of which improved editing outcomes in mammalian systems, revealed no significant improvement in gene editing in protoplast systems. This result may be explained by fundamental biological differences in cellular uptake mechanisms and intracellular environments, as well as differences in transfection and treatment methods. This latter difference includes the use of PEG transfection instead of the lipofection or electroporation methods used for mammalian cell transfection, in addition to differences in media pH and salt concentration. While PEG-free conditions did not produce detectable editing, future work should further explore applications of these RNP formulations for generating PEG-free editing in protoplasts, a desirable platform for maintaining cell viability and enhancing regeneration outcomes. While covalent attachment of cell penetrating peptides did not improve editing when using PEG transfection methods, optimization of this approach could be explored to generate PEG-free editing, where CPP amendments provide an alternate route for RNP internalization. Testing such co-incubation methods of RNPs and protoplasts at higher concentrations and for longer time periods may also yield enhanced editing.

In conclusion, the generation of stable polymeric RNP-NPs presents a simple and straightforward approach to improve gene editing efficiencies in plant protoplasts. All reagents are commercially available and require only a single additional preparation step relative to standard PEG transfection protocols. An outstanding question is whether anionic polymer stabilization is translatable to the other CRISPR/Cas protein orthologs that have been tested in plants. Additionally, in the future, this approach could be used in combination with other peptide-based methodologies, such as endosomolytic peptides, to expand the effectiveness of such tools in different species and plant tissue types.

## Methods

### Materials

All experiments were conducted with NLS-rich, *S. pyogenes* derived apo-Cas9 protein expressed and purified by the University of California Berkeley QB3 MacroLab. *S. pyogenes* Cas9 single guide RNAs (sgRNAs) were purchased from IDT with the manufacturer-recommended chemical modifications (Alt-R) and resuspended in IDT duplex buffer. Spacer sequences are listed in SI Table 1. Lyophilized polymers were purchased and prepared as previously described and stored at -80°C. Peptides with C-terminal Tyrosinse residues for Tyrosinase-mediated conjugation were purchased as lyophilized powder (GenScript) and resuspended to 1mM in water and stored at -30°C. The peptide A5K was purchased from CPC Scientific [SKU = IMMO-007A] and resuspended in DMSO to 10mM and stored at -30°C. Lyophilized Tyrosinase isolated from *Agaricus bisporus* (abTyr) was purchased from Sigma-Aldrich [SKU = T3824] and rehydrated to a working concentration of 2 mg/mL in 50mM phosphate buffer (pH = 6.5) and stored at -80°C.

### RNP-NP Preparation

Cas9 RNPs were freshly prepared day-of for all experiments reported. Stocks of sgRNA and apo-Cas9 were thawed on ice, diluted individually to 15μM and 10μM in NEBuffer 3 (100 mM NaCl, 50 mM Tris-HCl, 10 mM MgCl_2_ 1 mM DTT, pH = 7.9 at 25°C) respectively, and brought to room temperature. They were mixed 1:1 v/v to a molar ratio of 1:1.5 apoCas9:sgRNA and a target RNP concentration of 4 μM. When included, the PGA polymer was added at this step to a final concentration of 5 mg/mL (Nguyen et al. 2020). The RNP mixture with or without the polymer was incubated for 5 min at 37°C and then placed on ice. To prepare samples including the peptide A5K, we followed the previously described protocol, PERC. In brief, RNP was prepared as described above, and the peptide was diluted to 1mM at a 1/1/8 volume ratio of peptide / 2% Tween / nuclease free water. Where appropriate, water was used as the peptide solvent, however, for A5K this mixture contains residual DMSO necessary for suspension of the peptide at high concentrations. As relevant, particle size of RNP-NPs dispersed in NEBuffer 3 was measured by DLS on a Zetasizer Nano (Malvern Panalytical).

### In vitro Cleavage Assay

Cas9 RNPs were assembled as described above and diluted to 400 nM. DNA oligonucleotide containing the target cut site was generated by PCR amplification, purified, and stored at a concentration of 100 ng/uL at -20°C in water. Cleavage reactions were initiated by combining RNP with DNA at a molar ratio of 5:1 with a final DNA concentration of 5 ng/μL in either NEBuffer3 or the Transfection Buffer, then incubated at 37°C in a thermocycler for 16 hr. The enzyme was inactivated by holding the reaction mixture at 80°C for 15 minutes, then the reaction mixture was run on a 1.5% agarose gel containing TBE pre-stained with SYBR Safe (ThermoFisher Scientific) and a Quick-Load 100 bp DNA Ladder (New England Biosciences).

### Cas9-peptide tyrosinase-mediated conjugation

Coupling of Cas9 to peptides followed previously published methods. In brief, fresh aliquots of 40 μM apo-Cas9 were buffer-exchanged to remove TCEP using Zebra Spin Desalting Columns with a 7K MWCO (ThermoFisher Scientific). Peptide in five-fold molar excess and tyrosinase were combined with apo-Cas9 at 15 μM for 1 hr at 4°C. Samples were quenched with 2 mM tropolone for five minutes, then solvent exchanged into buffer containing 20 mM Tris HCl, 75 mM KCl, 5 mM MgCl_2_, 5 mM TCEP, 5% glycerol, pH = 7.5 using the same type of desalting column described above.

For samples to be analyzed, we performed an additional desalting step to remove all residual salts, then added formic acid to 5% by volume. We performed electrospray ionization mass spectrometry (ESI-MS) of the conjugate products using an Agilent 1260 series liquid chromatography outfitted with an Agilent 6224 time-of-flight (TOF) LC-MS system (Santa Clara, CA) courtesy of the Francis Lab at UC Berkeley. We employed a “shotgun” approach which bypassed the analytical column and delivered our product directly to the detector. Data was collected and analyzed by deconvolution using Agilent MassHunter Qualitative Analysis software. The open source Chartograph software (www.chartograph.com) was used for plotting the MS traces.

### Protoplast isolation and Transfection

Protoplasts were isolated following the previously reported tape-sandwich method (Wu et al. 2009). Masking tape (Duck band) was used to cover the adaxial surface of the leaf, and clear packing tape (Uline brand), was used to remove the abaxial epidermis before enzymatic digestion. For protoplast generation from *N. benthamiana*, 2.5-3.5 week old plants were used. Digestion solution was prepared using 20 mM MES (pH 5.7) 0.4 M mannitol and 20 mM KCl. The solution was warmed to 55°C before addition of 1.5% (wt/vol) cellulase R10 and 0.4% (wt/vol) macerozyme R10. Once cooled to room temperature, 10 mM CaCl2 and 0.1% BSA were added. Covered cells were digested with gentle shaking (40 rpm) for 2.5 hours at 30°C followed by 0.5 hours at 37°C. Digestion of 6-7 week-old *A. thaliana* was carried out using a solution prepared in the same way but with 1% (wt/vol) cellulase and 0.25% (wt/vol) macerozyme. Digestion time was 2 hours at 30°C. Subsequent wash and transfection steps were conducted using previously described buffer compositions (Yoo et al. 2007). Cells were filtered through a 0.45 μm cell strainer and centrifuged for 3 minutes at 100 rcf in round bottom culture tubes with ramping speeds adjusted to 0. Cells were washed with W5 buffer and centrifuged with the same conditions. After a second wash with W5, cells were allowed to settle on ice in W5 for 30 minutes before removal of W5 buffer and replacement with MMG buffer for transfection. Cells were counted using a hemocytometer and diluted to 1*10^6 cells/mL in MMG. For PEG transfection of *N. benthamiana* protoplasts, 20 μL of RNPs formulated as described above were first mixed with 100 μL of cells, followed by the addition of 110 μL of PEG solution. These volumes were doubled for *A. thaliana* protoplasts to guarantee sufficient genomic DNA yields. After a 5 minute transfection period, cells were diluted with 0.5 mL W5 and centrifuged 100 rcf for 3 minutes. The supernatant was then removed, and protoplasts were resuspended in 1 mL of WI buffer and moved to 12-well plates. Transfected protoplasts were kept covered in foil at 26°C for 24 hours post-transfection, at which point they were pelleted and frozen in liquid nitrogen.

### Amplicon Sequencing and Analysis

Genomic DNA was extracted from treated protoplasts derived from *N. benthamiana* and *A. thaliana* using Qiagen DNeasy plant mini kit (Qiagen 69106). 200-300bp regions containing the respective target cut sites were amplified and labeled with common primer overhangs using 30-cycle PCR with the primers listed in SI Table 2. These amplicons were purified using 1 x Ampure XP beads (UC Berkeley, DNA Sequencing Facility) and ran on a 2% agarose gel to confirm size. These samples were then sent to the IGI Next Generation Sequencing Core for a second round of PCR and subsequent sequencing using an Illumina MiSeq Kit v3 (600 cycles) with a read length of 2 x 300 bp. Reads were processed using CRISPResso2 software with default settings unless otherwise specified. To generate the percent intended editing efficiencies reported, we defined the following parameters: quantification window center = -3; quantification window size = 10; number of processes = 4; ignore substitutions. With these parameters, the resulting quantification values only include insertions and deletions (indels). Unless otherwise stated, all values reported are the averages of three technical replicates. Prism software (GraphPad) was used for statistical analysis and visualization. For Fig 2a and 2b, an unpaired T-test was used to determine statistical significance. Results shown in SI Fig. 2 were sequenced at an earlier date at the QB3 Genomics core using their Illumina MiSeq v3 instrumentation. It was confirmed that results were consistent when sequencing the same sample across the two different cores.

### Protoplast Viability Assay

Fluorescein diacetate (FDA) was diluted in acetone to a stock concentration of 5 mg/mL. Propidium iodide (PI) was diluted in water to a stock concentration of 1 mg/mL. Cells were diluted 10-fold from transfection concentrations in WI buffer to a final volume of 100 μL in a 48-well plate, and stains were added to final concentrations of 0.1 mg//mL for FDA and 2.5 μg/mL for PI. Cells were incubated for 20 minutes in dark conditions before imaging with the ImageXpress Micro cellular imaging system at 10x magnification with Cy5, TxRed, and GFP filters. Each transfection condition was repeated in triplicate, and eight images with each filter were taken within each well, corresponding to 24 images per channel per condition. Staining and imaging was conducted at 0hr and 24hr post transfection time points. Cell counting was conducted using the Cell Scoring Application Module of the MetaXpress software. Cells were identified using chloroplast autofluorescence detected in the Cy5 channel. Image sites with fewer than 50 cells in the field of view were excluded from analysis. Dead cells were those containing PI nuclear signal detected using the TxRed channel, and alive cells were those containing FDA cytoplasmic signal detected using the GFP channel. The percentage of dead or alive cells was calculated by dividing total number of cells (Cy5 positive) by total number of dead (PI positive) or alive (FDA positive) cells.

## Author Contributions

All authors contributed to the study conception and design. JH and ST contributed equally to this work, including conceptualization, material preparation, data collection and analysis, and manuscript generation. NC, DH, and TY contributed to experimental work. JM contributed to project conceptualization and experimental work. ML contributed to conceptualization and manuscript preparation.

## Data Availability

The datasets generated during and/or analysed during the current study are available in the NCBI SRA repository under BioProject PRJNA1197535

## Acknowledgements

We thank C. Jeans and the QB3 MacroLab at UC Berkeley for providing Cas9-NLS purified protein for RNP experiments. We thank M. Francis and the Francis Lab at UC Berkeley for guidance on and instrumentation use for covalent conjugation and QTOF methods. We thank M. West of the Cell and Tissue Analysis Facility (CTAF) at UC Berkeley. This work was performed in part in the QB3 CTAF, that provided the ImageXpress Micro cellular imaging system and MetaXpress software and training on each platform for automated cell imaging. We thank N. Krishnappa and the Innovative Genomics Institute Next Generation Sequencing Core for providing sequencing via Illumina MiSeq Kit v3. We thank QB3 Genomics, UC Berkeley, Berkeley, CA, RRID:SCR_022170 for assistance with sequencing of select samples via Illumina MiSeq v3.

