## supplemental information for "Anionic polymers stabilize Cas9 ribonucleoprotein nanoparticles to improve direct protein delivery and genome editing efficiencies in plant protoplasts"

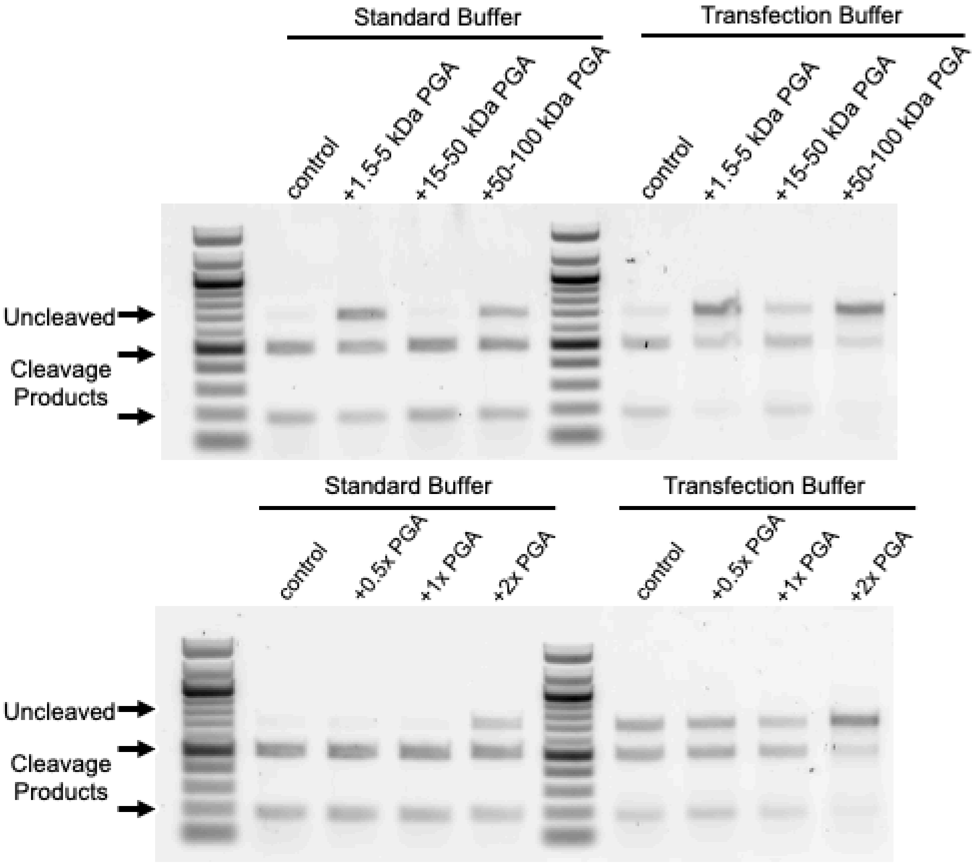


**Supplementary Fig. 1 |** *In vitro* cleavage assay run to optimize PGA formulation for polymer size and concentration.

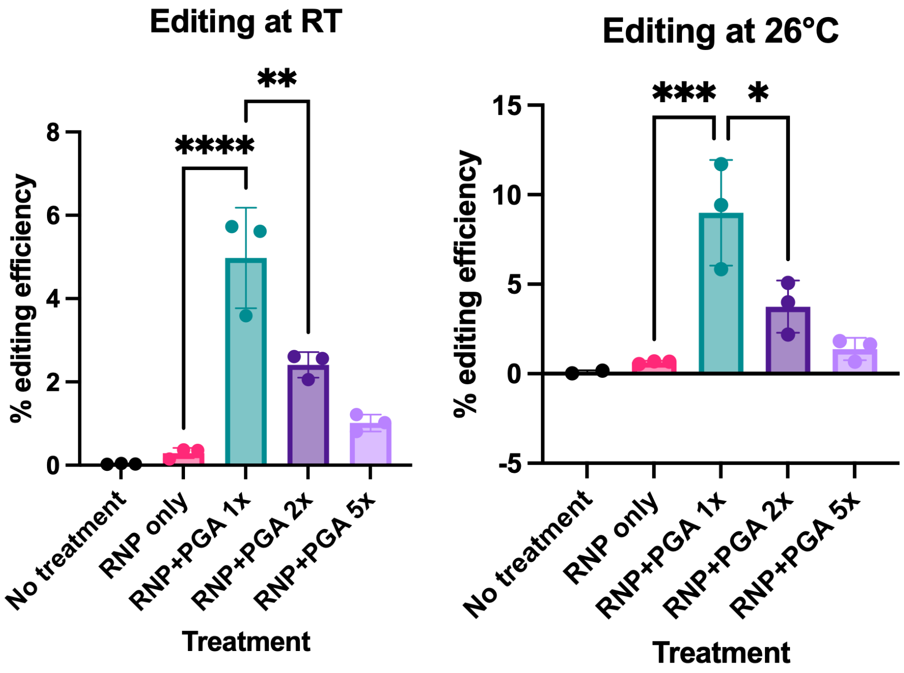


**Supplementary Fig. 2 |** Gene editing efficiencies of PEG-transfected Cas9 RNPs to protoplasts that were **(a)** formulated with or without the anionic polymer PGA and sgRNA1 to *N. benthamiana* protoplasts. For all editing results, editing efficiencies were quantified using CRISPResso2 software to analyze NGS results. Mean and SD of three technical replicates are shown. ANOVA test was used to calculate *P* value of indicated comparisons. $*P \leq 0.05$, $**P \leq 0.01$, $***P \leq 0.001$, $**** P \leq 0.0001$. Generated using QB3 Genomics core data.

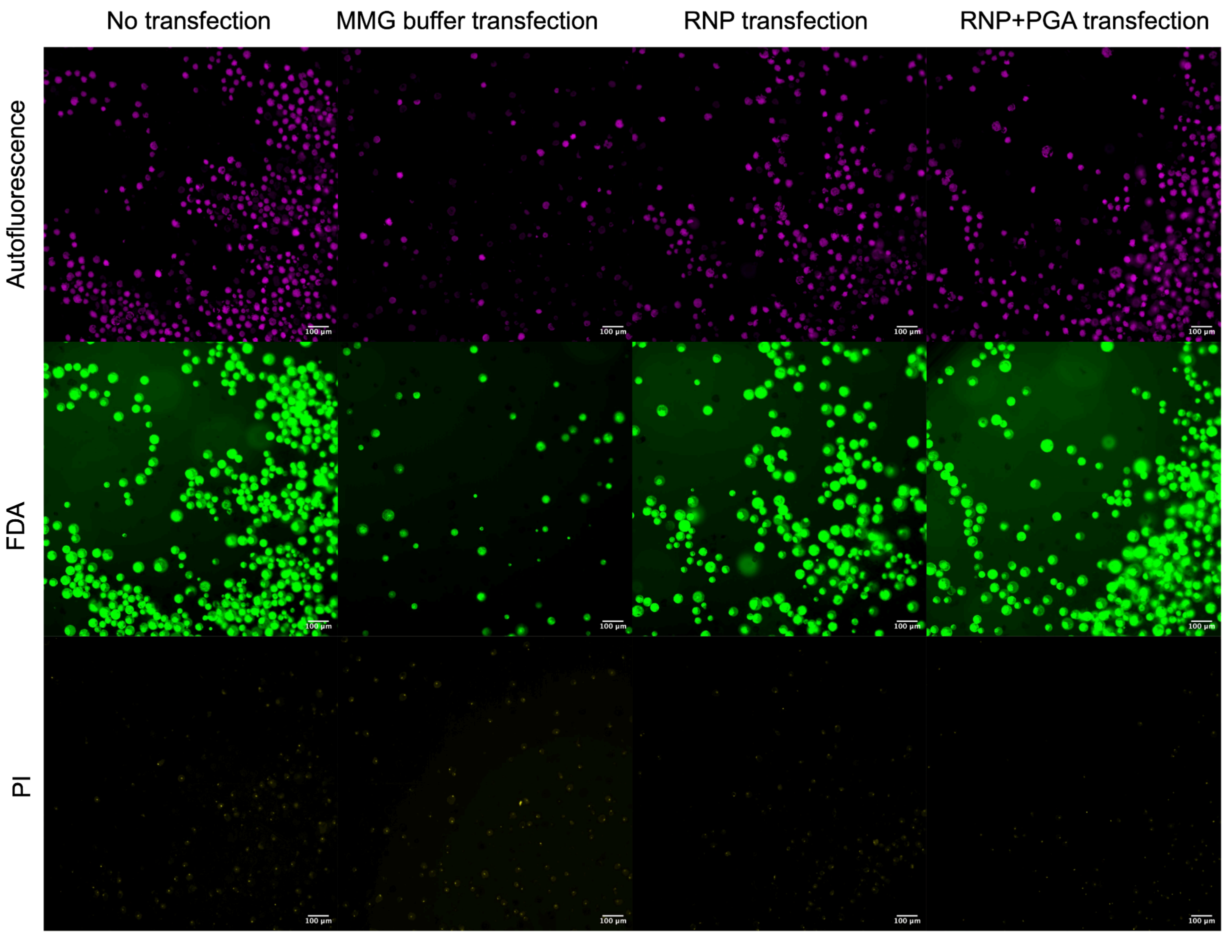


**Supplementary Fig. 3 |** Representative images for viability screening, shown for un-transfected protoplasts at 24hr post-treatment. Viability was screened with FDA and PI dual staining using the IXM cellular imaging system. Chlorophyll autofluorescence (magenta) indicates individual protoplasts, PI staining (yellow) in the nucleus indicates a dead cell, and FDA staining (green) in the cytoplasm indicates a viable cell.

| | Peptide Sequence | Peptide MW (Da) | Approximate Predicted Conjugate Product MW (Da) |
|---|---|---|---|
| Cas9 | - | - | 161000 |
| Cas9-2xFLG22 | QRLSTGSRINSAKDDAAGLQIAGGY | 2500 | 166000 |
| Cas9-2xR9 | RRRRRRRRRGGY | 1700 | 164400 |
| Cas9-2xSV40 | KKKRKVGGSPKKKRKVGGY | 2100 | 165200 |
| Cas9-2xWUS | KNVFYWFQNHKARERQKKRFNGGY | 3100 | 167200 |

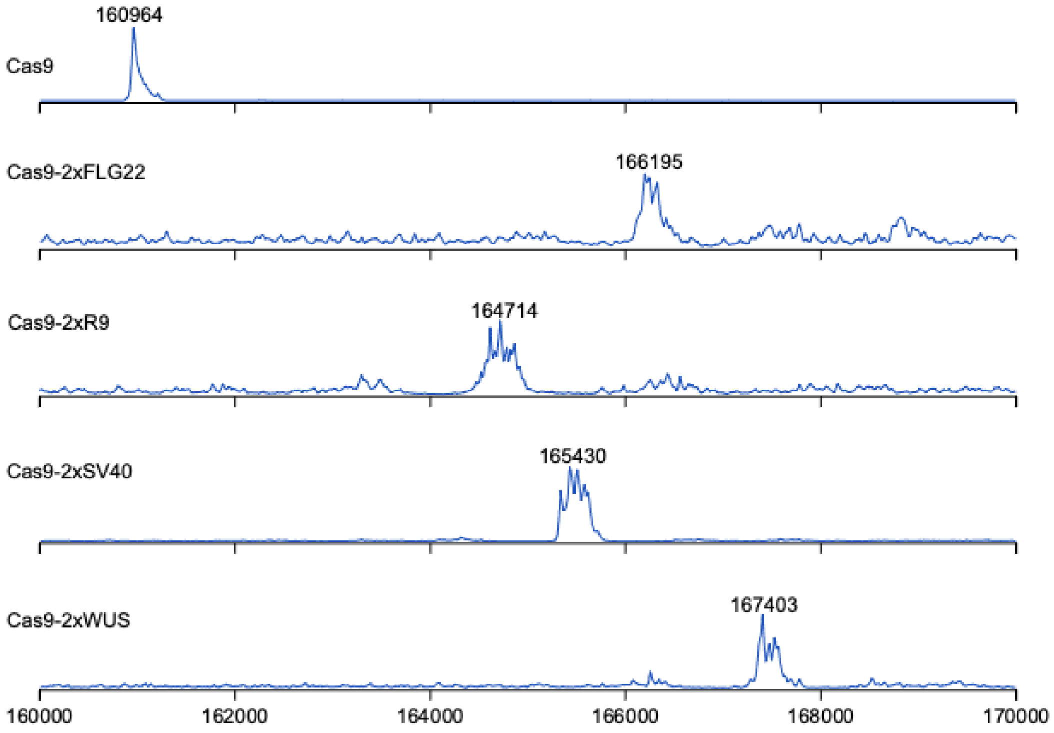


**Supplementary Fig. 4 |** Conjugation results QTOF.

| Name | Gene Target | Spacer sequence [PAM] | Plant Species | Reference |
|---|---|---|---|---|
| sgRNA1 | NbPDS | GCCGTTAATTTGAGAGTCCA [AGG] | Nicotiana benthamiana | Nekrasov, 2013 |
| sgRNA2 | NbPDS | TTGGTAGTAGCGACTCCATG [GGG] | Nicotiana benthamiana | Uranga, 2023 |
| sgRNA3 | AtPDS3 | GGACTTTTGCCAGCCATGGT [CGG] | Arabidopsis thaliana | Li, 2013 |

**Supplementary Table 1 |** Guide RNAs used in this study.

| Name | Forward Primer Sequence | Reverse Primer Sequence | Assay |
|---|---|---|---|
| sgRNA1 | TGCATAGTATTTAGGTTCACAAGTGGG | TCCTTTGTCAATCTTCGGGTCGT | NGS |
| sgRNA2 | TCAATAAAATGCCCCAAATTGGACTTG | TCTTATTTTAGAGGATATATGCCCACGATCATAAATC | NGS |
| sgRNA3 | GTTGTTGCTGTTGGATTTACG | CACAACAACCACATGGACTAG | NGS |

**Supplementary Table 2 |** Oligonucleotides used for generating amplicons that were analyzed by NGS.
